# Genetic compensation of triacylglycerol biosynthesis in the green microalga *Chlamydomonas reinhardtii*

**DOI:** 10.1101/2021.08.12.455529

**Authors:** Yi-Ying Lee, Rudolph Park, Stephen M. Miller, Yantao Li

## Abstract

Genetic compensation through transcriptional adaptation has been proposed to explain phenotypic differences between gene knockouts and knockdowns. With the rapid development of reverse genetic tools such as CRISPR/Cas9 and RNAi in microalgae, it is increasingly important to assess whether genetic compensation affects the phenotype of engineered algal mutants. While exploring triacylglycerol (TAG) biosynthesis pathways in *Chlamydomonas reinhardtii*, it was discovered that knockout of certain genes catalyzing rate-limiting steps of TAG biosynthesis, type-2 diacylglycerol acyltransferase genes (*DGTTs*), triggered genetic compensation under abiotic stress conditions. Genetic compensation of a *DGTT1* null mutation by a related *PDAT* gene was observed regardless of the strain background or mutagenesis approach, e.g., CRISPR/Cas 9 or insertional mutagenesis. However, no compensation was found in the *PDAT* knockout mutant. The effect of *PDAT* knockout was evaluated in a Δ*vtc1* mutant, in which *PDAT* was up-regulated under stress. Knockout of *PDAT* in the Δ*vtc1* background induced a 12.8-fold upregulation of *DGTT1* and a 272.3% increase in TAG content in Δ*vtc1*/*pdat1* cells, while remaining viable. These data suggest that genetic compensation contributes to the genetic robustness of microalgal TAG biosynthetic pathways, maintaining lipid and redox homeostasis in the knockout mutants under abiotic stress with a mechanism distinct from that found in plants. This work demonstrates examples of genetic compensation in microalgae, implies the physiological relevance of genetic compensation in TAG biosynthesis, and provides guidance for future genetic engineering and mutant characterization efforts.

## Introduction

Organisms have evolved various genetic buffering systems to maintain fitness in response to genetic perturbations, including functionally redundant genes (Tautz, 1992), protein feedback loops (Barabási & Oltvai, 2004), and acquisition of adaptive mutations (Teng et al., 2013). Such buffering or compensation systems ensure similar growth or developmental outcomes despite some genetic changes (El-Brolosy & Stainier, 2017). Genetic compensation is defined as the phenomenon whereby the effect of a deleterious mutation is buffered by the genome (Kontarakis & Stainier, 2020). Recently, genetic compensation through transciptional adaptation, by which a homologous gene or genes are upregulated in response to an early nonsense mutation in a related gene, has been proposed as a mechanism for maintaining genetic robustness in a number of metazoans, including zebrafish, mice, and nematodes, and to explain profound differences between gene knockouts and knockdowns at certain loci (El-Brolosy et al., 2019; Z. Ma et al., 2019; Rossi et al., 2015; Serobyan et al., 2020). Likewise, comparison between gene knockouts and knockdowns in some model plant systems has revealed puzzling discrepencies that suggest transcriptional adaptation is acting to buffer the genome in these species as well (Braun et al., 2008; Chen et al., 2014; Gao et al., 2015; Rodriguez-Leal et al., 2019).

Increasingly, gene knockdown and knockout tools have been deployed in the green microalga *Chlamydomonas reinhardtii* (Chlamydomonas; (Greiner et al., 2017; Jiang, Brueggeman, Horken, Plucinak, & Weeks, 2014; Xiaobo Li et al., 2019; Molnar et al., 2009; Rohr, Sarkar, Balenger, Jeong, & Cerutti, 2004; Schroda, 2006)), but little attention has been paid as to whether these tools might for some genes lead to very different outcomes, and more broadly, whether genetic compensation occurs in green algae (El-Brolosy & Stainier, 2017; Kontarakis & Stainier, 2020). Here we set out to address these questions as part of an ongoing study of lipid biosynthesis in Chlamydomonas, using triacylglycerol (TAG) biosynthesis pathways as examples.

Microalgae store TAG as a carbon and energy-rich compound in cells, whose synthesis can be stimulated under various stress conditions (Hu et al., 2008; Lenka et al., 2016). Nutrient deprivation such as nitrogen (N) deprivation and phosphorus (P) deprivation are most commonly used stressors to induce TAG production. Under N deprivation, algal cell growth is almost completely arrested while P deprivation promotes TAG production with moderate cell growth (Iwai, Ikeda, Shimojima, & Ohta, 2014). Biosynthesis of TAG involves sequential acylation to the precursor glycerol-3-phosphate. Two pathways are involved in the last step of TAG biosynthesis. One pathway is a *de novo* acyl-CoA-dependent route mediated by diacylglycerol acyltransferase (DGAT) (Ohlrogge & Browse, 1995). Three major types of DGATs including two structurally distinctive, membrane bound type-1 and type-2 DGATs and a soluble cytosolic type-3 DGAT have been identified in plants, algae, and other eukaryotes (Cases et al., 1998; Lardizabal et al., 2001; Oelkers, Cromley, Padamsee, Billheimer, & Sturley, 2002). The other TAG biosynthesis pathway is an acyl-CoA-independent route that requires membrane lipid breakdown and is mediated by a plastidic membrane bound enzyme, phospholipid:diacylglycerol acyltransferase (PDAT) (Dahlqvist et al., 2000; Yoon, Han, Li, Sommerfeld, & Hu, 2012).

The genome of *C. reinhardtii* harbors one type-1 *DGAT* gene (*DGAT1*), five type-2 *DGAT* genes (*DGTT1-5*), and one *PDAT* gene (Bagnato et al., 2017; Boyle et al., 2012; Miller et al., 2010; Yoon et al., 2012). RNA-Seq analysis previously revealed that the abundance of *PDAT*, *DGAT1* and *DGTT1* transcripts was up-regulated, and that of *DGTT2* and *DGTT3* was moderate but constant, and the abundance of transcripts of *DGTT4* and *DGTT5* was low to undetectable in response to N deprivation (Boyle et al., 2012). Subsequent RT-qPCR analyses corroborated these findings for *PDAT*, *DGAT1* and *DGTT1*, while *DGTT2-3* were found to be induced by N deprivation (Liu, Han, Yoon, Hu, & Li, 2016; Yoon et al., 2012). Similar results were found in response to P deprivation, where *PDAT*, *DGAT1*, and *DGTT1-4* were up-regulated during TAG accumulation (Iwai et al., 2014). Functional studies using *in vitro* enzyme assays and gene knockdowns have largely yielded results consistent with expectations based on these expression data. *DGTT1-3* and *PDAT* were shown to catalyze TAG biosynthesis *in vitro* (Liu et al., 2016; Yoon et al., 2012). Knockdown of *DGTT1*, *DGTT2* or *DGTT3* each resulted in a moderately reduced TAG (TAG-less) phenotype under persistent stress, while knockdown of *PDAT* resulting in a moderate TAG-less phenotype during both favorable growth and the early phase of stress induction (Liu et al., 2016; Yoon et al., 2012), suggesting *DGTT*s and *PDAT* are functionally related genes involved in TAG biosynthesis in Chlamydomonas. However, in some cases, analysis of gene knockdowns can be limited by off-target effects and low efficiency of gene silencing (Fedorov et al., 2006; Y. Ma, Creanga, Lum, & Beachy, 2006), so null mutants are needed to better understand gene function. Among Chlamydomonas TAG biosynthesis genes, so far only *pdat* mutants have been characterized, which had reduced TAG content under stress (Boyle et al., 2012). However, whether the *PDAT* insertion mutations analyzed were buffered by expression of related genes such as *DGTT*s is not known, and up to now no *DGTT* mutants have been analyzed. So the precise contributions of these genes to TAG accumulation remains obscure.

In this work we analyzed TAG accumulation in strains bearing null mutations in several genes including *DGTT1-3* and *PDAT*, and found phenotypes that contradicted known knockdown phenotypes in these genes. Our data showed genetic compensation through up-regulation of related TAG biosynthetic genes for *dgtt1* and *dgtt2* mutants resulted in a higher TAG content under stress, which is contrary to the TAG-less phenotype in the respective knockdown lines. Interestingly, no genetic compensation was found in the *pdat* mutant. However, when evaluated in the Δ*vtc1* background (defective for Vacuolar Transporter Chaperone 1, a component of the polyphosphate polymerase complex), where *PDAT* expression and function is likely critical for storing excess energy as TAG under stress, the *PDAT* knockout resulted in compensation by up-regulation of *DGTT1* and a higher TAG content. Our data demonstrate genetic compensation exists in microalgae and imply the physiological importance of genetic compensation in TAG biosynthesis under stress.

## Results

### Genetic compensation is present in microalgae

Type-2 DGATs are believed to be major contributors for TAG biosynthesis in algae, but previous knockdown of Chlamydomonas type-2 *DGAT* genes *DGTT1*, *DGTT2* and *DGTT3* caused only relatively small (~10 - 20%) suppression of TAG accumulation (Liu et al., 2016). We set out to determine whether null mutations in these genes might have larger effects, so we obtained *dgtt1*, *dgtt2* and *dgtt3* CLiP library knockout mutants generated by the Jonikas group (Xiaobo Li et al., 2019; Xiaobo Li et al., 2016) and analyzed their TAG accumulation. Each mutant strain carries a CIB1 cassette insertion that disrupts the respective *DGTT* gene. We further examined the genotype of these mutants and found the *dgtt1* mutant (LMJ.RY0402.223444) insertion is in the middle of intron 1, the *dgtt2* mutant (LMJ.RY0402.213587) insertion is at the junction between exon 4 and intron 4, and the *dgtt3* mutant (LMJ.RY0402.223565) insertion is at the beginning of exon 3 (Fig. 1A; Fig. S1). No detectable transcripts of *DGTT1*, *DGTT2* and *DGTT3* were found in the respective mutants (Fig. 1C), suggesting all the mutants are null.

**Fig 1.**
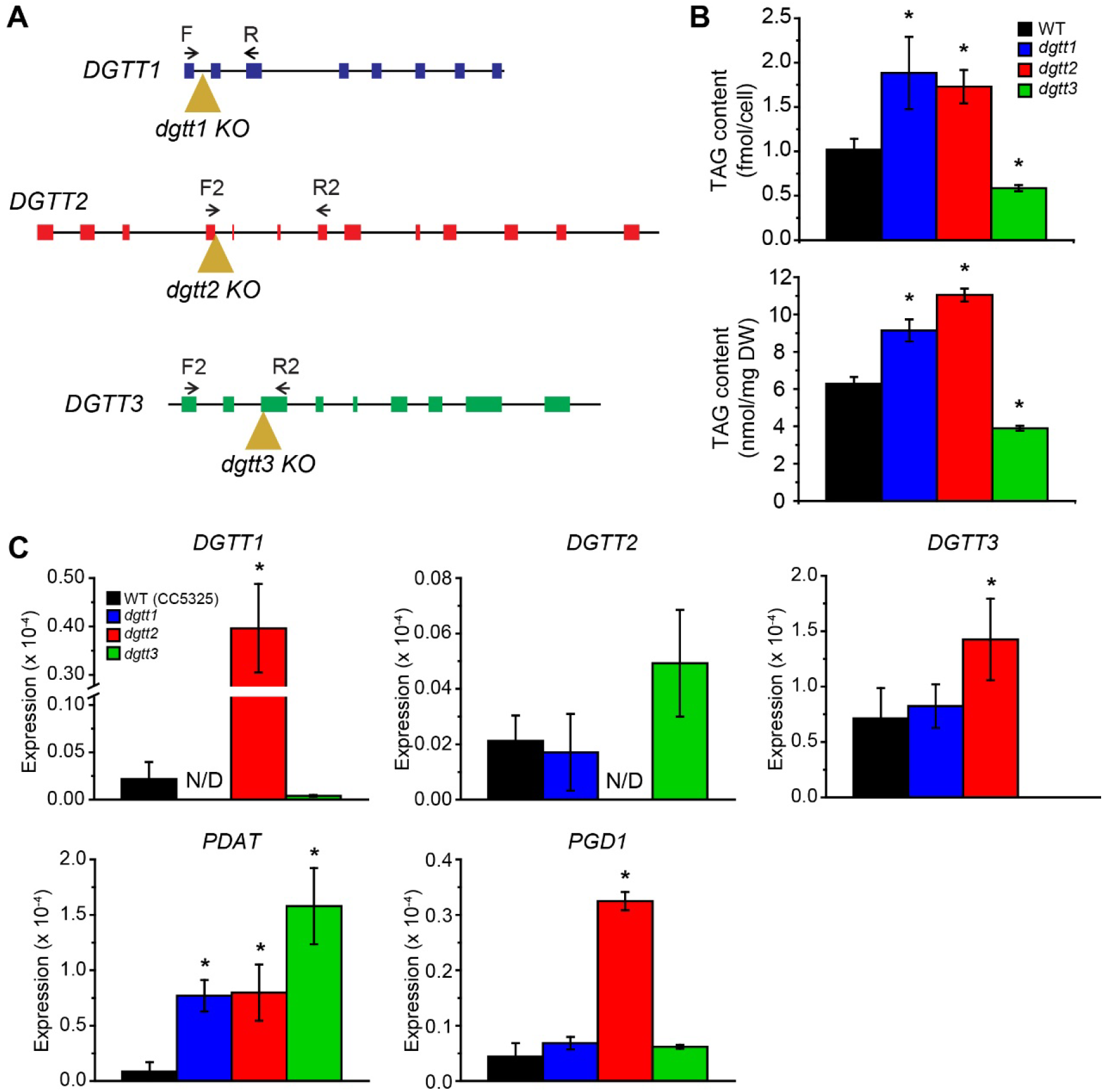
Genetic structure, phenotype and gene expression of the *dgtt1*, *dgtt2*, *dgtt3* knockout mutants obtained from the CLiP library. (A) Locations of the insertion sites of the CIB1 cassette (brown triangles) and of the primer pairs (black arrows) used to detect gene expression in the *DGTT1*, *DGTT2* and *DGTT3* genes are presented. (B) TAG content of the *dgtt1* (blue), *dgtt2* (red), *dgtt3* (green) mutants and the parent strain CC5325 (black, WT) grown was measured at 72 h under P deprivation. (C) Relative transcriptional level of selected TAG biosynthesis genes (*DGTT1*, *DGTT2*, *DGTT3*, *PDAT* and *PGD1*) in the *dgtt1* (blue), *dgtt2* (red), *dgtt3* (green) mutants and the parent strain CC5325 (black) was measured at 24 h under P deprivation. Transcripts of *DGTT1* in the *dgtt1* mutant and *DGTT2* in the *dgtt2* mutant were nondetectable (N/D). Data represents mean ± standard deviation (SD) from three biological repeats. An asterisk indicates statistical significance by Student’s *t*-test (*p* value ≤ 0.05) when compared to WT.

TAG biosynthesis was comparatively studied in the *dgtt1*, *dgtt2*, and *dgtt3* mutants and their parent strain (CC5325; wild-type) under P deprivation, which was reported to induce TAG accumulation in *C. reinhardtii* while maintaining moderate cell growth (Molnar et al., 2009). Surprisingly, the *dgtt1* and the *dgtt2* knockout mutants had significantly higher TAG content than the wild-type strain (t-test, p ≤ 0.05, Fig. 1B) either on a per cell basis or per dry weight (DW) basis. The TAG content of *dgtt1* and *dgtt2* mutant cells increased by about 90% and 70% per cell, respectively, and by 45% and 76% per unit dry weight, compared with wild-type cells at 72 h under P deprivation (Fig. 1B). These results are in stark contrast to those for the knockdown strains, which had reduced TAG content (Liu et al., 2016), and they suggest a compensation mechanism exists in Chlamydomonas that affects other TAG genes/enzymes in the *dgtt1* and *dgtt2* mutant. By contrast, the *dgtt3* knockout mutant contained ~40% less TAG per cell and per dry weight compared with the wild-type (Fig. 1B), and notably it had about twice the decrease in TAG as that caused by the *dgtt3* knockdown (Liu et al., 2016). In Chlamydomonas, *DGTTs* and *PDAT* are homologs. Sequence analysis revealed identities between *PDAT* and *DGTT1*, *DGTT2*, and *DGTT3* were 50.9%, 49.8%, and 49.2%, respectively (Table S1), all above 40% sequence identity needed to induce genetic compensation by homologs in previous studies (Z. Ma et al., 2019).

To explore possible genetic compensation in the knockout mutants, the expression of TAG biosynthetic genes was examined, including that of *DGTT1*, *DGTT2*, *DGTT3*, *PDAT*, and *PGD1* (encodes an MGDG-specific lipase and presumably functions with PDAT in membrane recycling) (X. Li et al., 2012; Liu et al., 2016; Yoon et al., 2012). In the *dgtt1* mutant, the expression of *DGTT2*, *DGTT3* and *PGD1* remained unchanged, but the expression of *PDAT* was upregulated by 9-fold compared with the wild-type at 24 h under P deprivation (Fig. 1C). In the *dgtt2* mutant, the expression of all tested genes was upregulated; compared with the wild-type, the expression of *DGTT1*, *DGTT3*, *PDAT* and *PGD1* in the *dgtt2* mutant was increased by 19-fold, 2.6-fold, 9-fold, and 7.6-fold, respectively, at 24 h under P deprivation (Fig. 1C), correlating with higher TAG accumulation in the *dgtt2* mutant. In the *dgtt3* knockout strain, the expression level of *PDAT* was significantly higher than for the wild type (t-test, p ≤ 0.05; Fig. 1C), but that increase did not lead to a functional compensation for TAG accumulation (Fig. 1B).

Our findings suggest that a mutation leading to a premature stop codon in two different type-2 *DGAT* genes likely activates a genetic compensatory network by upregulation of homologous genes in Chlamydomonas. To determine whether this genetic compensation was specific to insertional mutants in the CLiP library genetic background, or whether it might be a more general phenomenon of null mutants with premature stop codons, we used CRISPR/Cas9 to generate a *dgtt1* mutant in a different strain background (*C. reinhardtii* CC3403). In the resulting mutant (*CC-dgtt1*), the *DGTT1* gene carries a 1229-bp insertion corresponding to pHS_SaCas9 sequence (Cas9 expression vector) near the beginning of exon 3 of the coding region (Fig. 2A and Fig. S2A) that results in premature stop codons (Fig. S2B) and disrupts expression of the *DGTT1* gene. The TAG content of *CC-dgtt1* was comparable to that of the wild-type on a per cell basis, and 66% higher than the wild-type per unit dry weight under P deprivation (Fig. 2B). Moreover, this phenotype was maintained under N deprivation (Fig. S3). Similar to the situation for the *dgtt1* mutant from the CLiP library, RT-qPCR analyses revealed that *PDAT* expression was upregulated by 6.3-fold in the *CC-dgtt1* mutant, while other TAG biosynthesis genes (*DGTT2*, *DGTT3* and *PGD1*) remained the same as in the wild-type (Fig. 2C). Taken together, genetic compensation of *DGTT* seems to occur in Chlamydomonas mutants regardless of the strain background or early-frameshift mutant allele.

**Fig 2.**
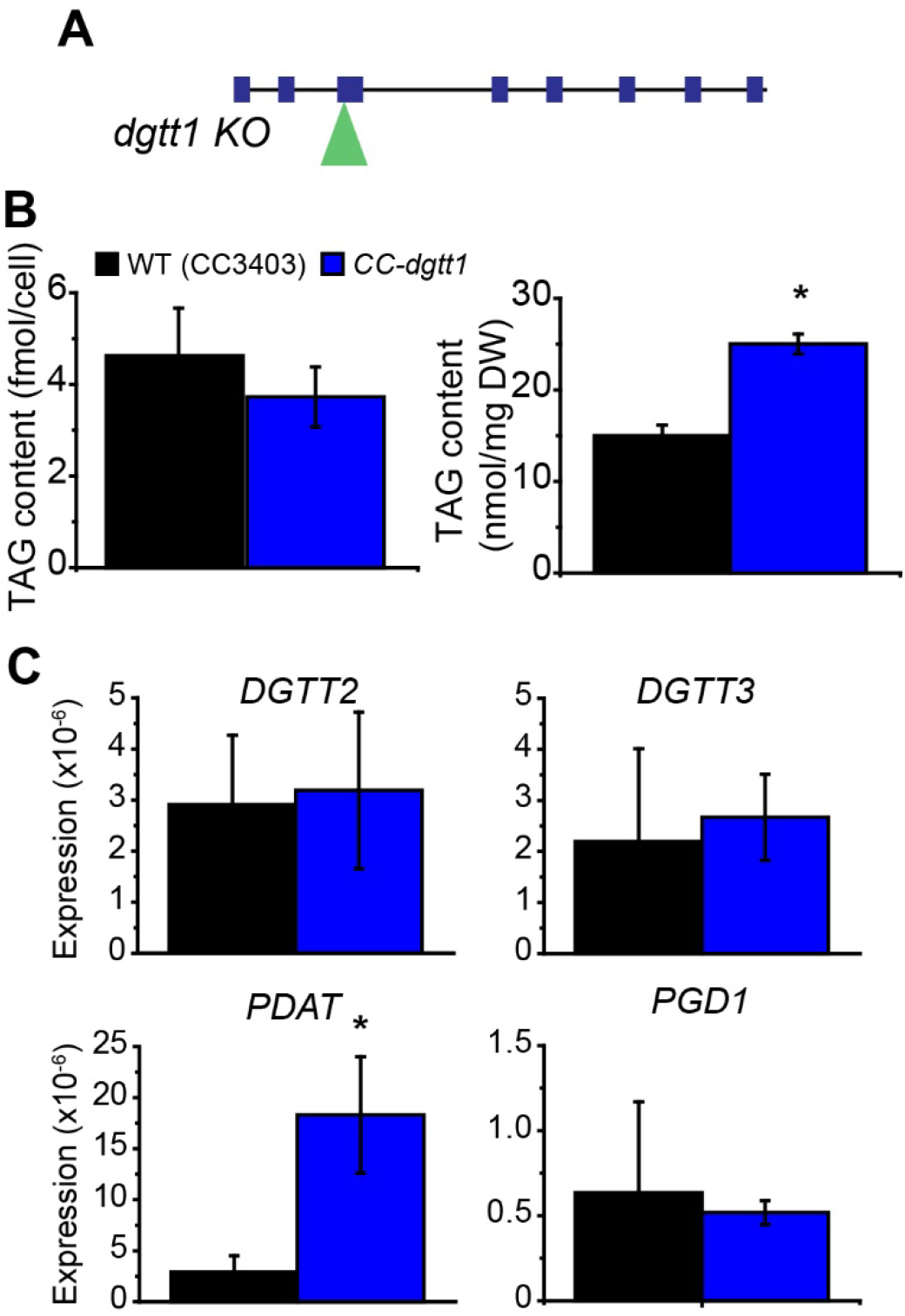
Genetic structure, phenotype and gene expression of the CRISPR-derived *dgtt1* knockout mutant. (A) The location of an *AphIII* cassette insertion (green triangle) generated by CRISPR in the *DGTT1* gene are presented. (B) TAG content of the *CC-dgtt1* mutant (blue) and the parent strain CC3403 (black, WT) grown was measured at 72 h under P deprivation. (C) Relative transcriptional level of selected TAG biosynthesis genes (*DGTT2*, *DGTT3*, *PDAT* and *PGD1*) in the *dgtt1* mutant (blue) and the parent strain CC3403 (black) was measured at 48 h under P deprivation. Data represents mean ± standard deviation (SD) from three biological repeats. An asterisk indicates statistical significance by Student’s *t*-test (*p* value ≤ 0.05).

### Compensation of *DGTT* knockouts is not reciprocal in the *pdat* mutant

Since *C. reinhardtii* cells are capable of compensating defects of *DGTT1* and *DGTT2* mutations by upregulating related genes such as *PDAT*, we asked whether the mechanism is reciprocal in the *pdat* mutant. To this end, we assessed TAG accumulation and the expression of the *DGTT1-3* genes in a *pdat* knockout mutant (CC4502) previously described in (Boyle et al., 2012). Unlike the *dgtt1 and dgtt2* mutants, the *pdat* mutant had reduced TAG levels (Fig. 3), similar to the *pdat* knockdown lines (Yoon et al., 2012). The relative TAG content of the *pdat* knockout mutant was ~30% lower than that of the wild-type strain at 48 h and 72 h under P deprivation (Fig. 3A), similar to the decrease in TAG content previously reported for this strain in response to N deprivation (Boyle et al., 2012). In P-deprived *pdat* cells, the expression of *DGTT1-3* were the same as or slightly lower than in the wild-type (Fig. 3B). These data suggest no genetic compensation of TAG biosynthesis occurs in the *pdat* mutant.

**Fig 3.**
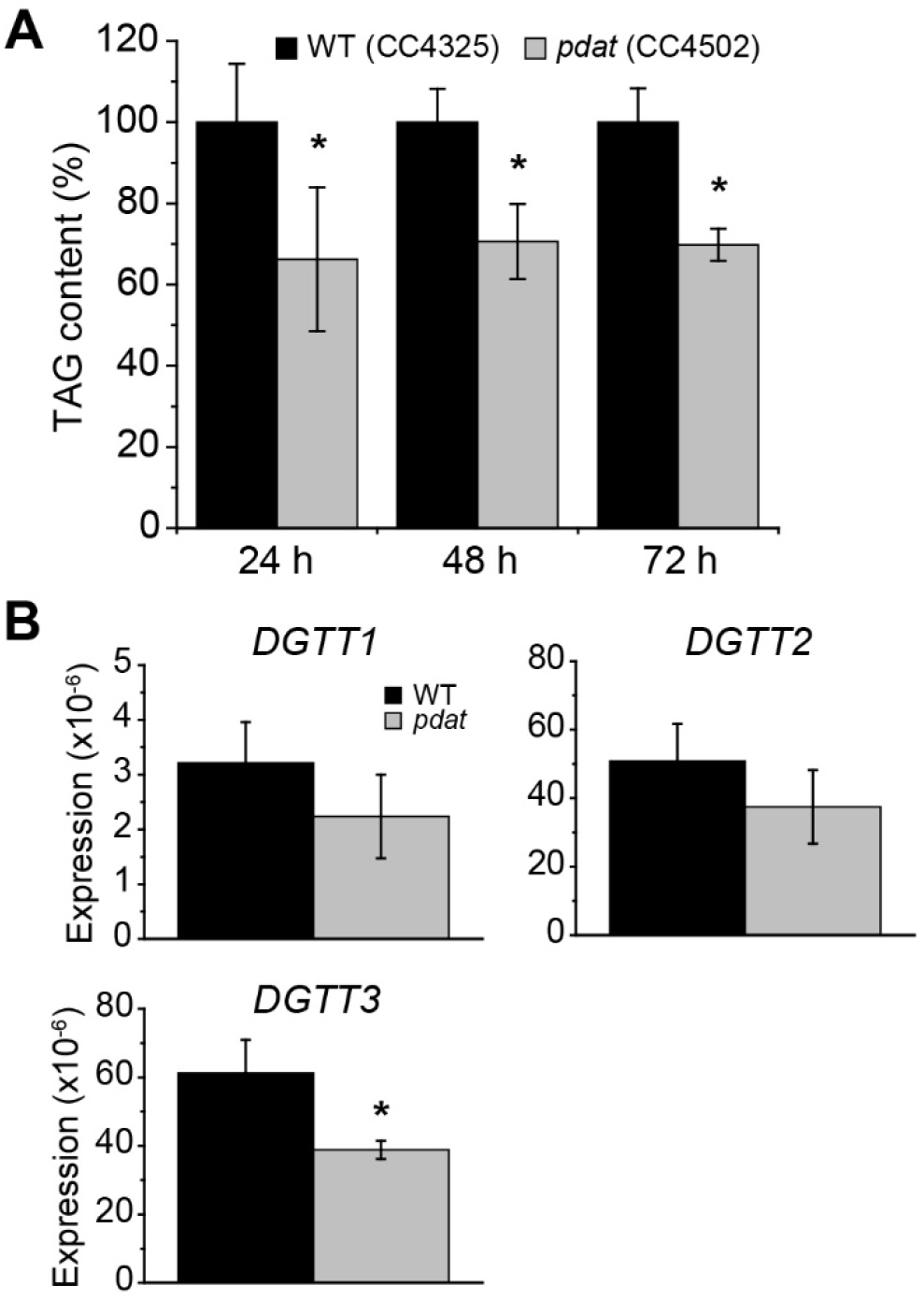
Phenotype and gene expression of the *pdat* mutant (CC4502). (A) Relative concentrations of TAG from the *pdat* knockout strain CC4502 (gray) and the wild-type CC4425 (black, WT) grown under 24 h-, 48 h-, 72 h- and 96 h-P deprivation was shown. (B) Relative transcriptional level of selected TAG biosynthesis genes (*DGTT1*, *DGTT2*, *DGTT3*) in the *pdat* mutant (gray) and the wild-type (black) was measured at 24 h under P deprivation. Data represents mean ± standard deviation (SD) from three biological repeats. An asterisk indicates statistical significance by Student’s *t*-test (*p* value ≤ 0.05).

### Compensation by upregulation of *DGTT1* in a Δ*vtc1/pdat1* double mutant

Our previous work suggests *PDAT* primarily regulates TAG biosynthesis under favorable, nutrient-replete conditions (Yoon et al., 2012) but not under persistent nutrient-deficient stress conditions (Liu et al., 2016). A possible explanation for lack of genetic compensation in the *pdat* mutant is thus the relatively poor function of *PDAT* under stress conditions, in which *DGTTs* are already highly expressed and responsible for TAG biosynthesis. Previous work showed P metabolism, particularly P uptake, was impaired in a Δ*vtc1* mutant, in which the vacuolar transport chaperon (VTC) complex was disrupted (Aksoy, Pootakham, & Grossman, 2014; Plouviez et al., 2021; Sanz-Luque, Saroussi, Huang, Akkawi, & Grossman, 2020; Schmollinger et al., 2021). As such, in this Δ*vtc1* mutant we reasoned that *PDAT* expression and function may be especially critical to recycle phospholipids and balance P metabolism under stress. To test this idea, we analyzed TAG biosynthesis in the Δ*vtc1* mutant and in a *VTC1*-rescued strain (*VTC1*) under P replete and P-depleted conditions. In Δ*vtc1* cells TAG content was 90% higher (*t*-test, p ≤ 0.05) than the control under P-deprived conditions but not under P-replete conditions (Fig. 4A and B), suggesting loss of *VTC1* had induced TAG biosynthesis under stress. When we analyzed the TAG biosynthetic genes in P-depleted Δ*vtc1* cells, only the *PDAT* gene showed an increase in expression (2.8-fold; Fig. 4C) while none of the *DGTT* or *PGD* genes were upregulated, suggesting a role of *PDAT* in controlling TAG biosynthesis under stress in the Δ*vtc1* mutant.

**Fig 4.**
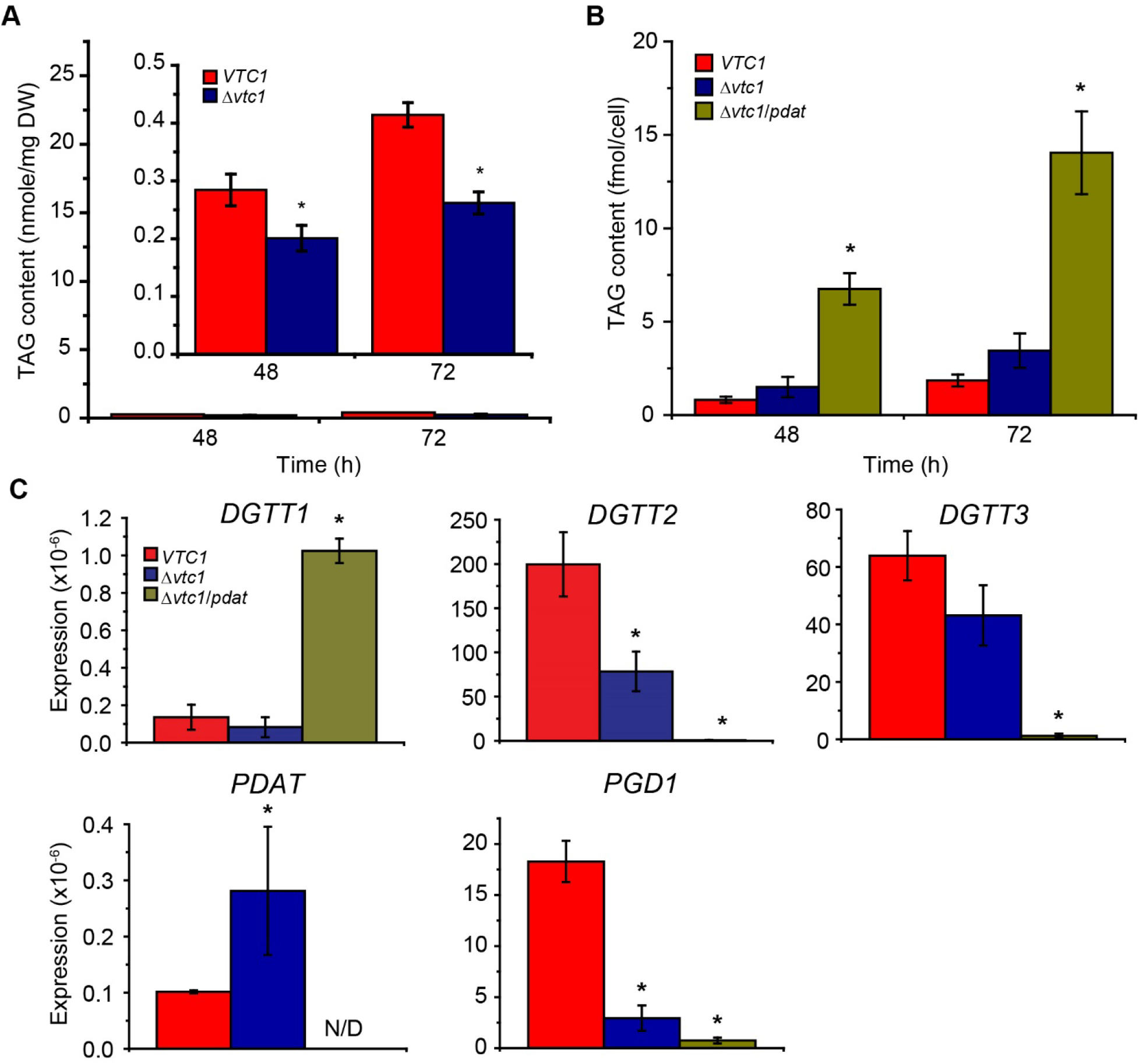
Phenotype and gene expression of the *VTC1* strain, Δ*vtc1* single mutant and Δ*vtc1*/*pdat* double mutant. (A) TAG content of the Δ*vtc1* strain CC5165 and the *VTC1* rescue strain CC5166 grown was measured at 48 h and 72 h under P replete. (B) TAG content of the *VTC1* strain, Δ*vtc1* strain and the Δ*vtc1*/*pdat* strain grown was measured at 48 h and 72 h under P deprivation. (C) Relative transcriptional level of selected genes (*DGTT1*, *DGTT2*, *DGTT3*, *PDAT* and *PGD1*) was measured at 24 h under P deprivation. Transcripts of *PDAT* in the Δ*vtc1*/*pdat* double mutant were nondetectable (N/D). Data represents mean ± standard deviation (SD) from three biological repeats. An asterisk indicates statistical significance by Student’s *t*-test (*p* value ≤ 0.05).

We predicted that for the Δ*vtc1* mutant, in which *PDAT* function is critical, if *PDAT* expression is interrupted, genetic compensation by other TAG biosynthetic genes may be necessary for the resulting mutant to survive under stress. To test this idea, we generated a Δ*vtc1*/*pdat* double mutant by crossing the Δ*vtc1* (CC5165) and the *pdat* (CC4502) single mutant strains. Interestingly, the TAG level in the resulting strain increased compared with the control and single mutant (Fig. 4B). After 48 h of P deprivation, the TAG content of Δ*vtc1*/*pdat* cells was about 272.3% higher than the *VTC1* control and 95% higher than the Δ*vtc1* mutant (Fig. 4B). We found that under stress, *DGTT1* transcripts were upregulated in Δ*vtc1*/*pdat* cells compared with the control (by12.4-fold), but no other TAG biosynthetic genes were transcriptionally upregulated (Fig. 4C). Moreover, Δ*vtc1*/*pdat* cells remained viable with moderate growth under P deprviation. Δ*vtc1*/*pdat* cell density slightly increased from 0.69 × 10^6^ cells/ml to 1.01 × 10^6^ cells/ml between day 0 and day 4 (Fig. S4A), while the per cell dry weight of the Δ*vtc1*/*pdat* mutant about doubled compared with the control (*VTC1*) (Fig. S4B). By contrast, the growth rate and per cell dry weight of Δ*vtc1* cells were similar to that of the control (Fig. S4). Therefore, our results suggest that upregulation of *PDAT* in the Δ*vtc1* mutant results in an increase in TAG accumulation, while the effect of *PDAT* disruption in the Δ*vtc1* background is buffered by upregulation of *DGTT1* in the *Δvtc1/pdat1* double mutant.

## Discussion

Genetic compensation, and more specifically, transcriptional adaptation through upregulation of related genes, is proposed as an underlying mechanism to explain discrepencies between knockdown and knockout phenotypes in several model systems including zebrafish and mouse (El-Brolosy et al., 2019; Z. Ma et al., 2019; Rossi et al., 2015), *Caenorhabditis elegans* (Serobyan et al., 2020), Arabidopsis (Braun et al., 2008; Chen et al., 2014; Gao et al., 2015), and tomato (Rodriguez-Leal et al., 2019). One mechanism for this phenomenon involves the nonsense-mediated mRNA decay pathway that is activated by early nonsense mutations (El-Brolosy et al., 2019; Z. Ma et al., 2019). However, it is not known whether this phenomenon exists across eukaryotes, including microalgae (El-Brolosy & Stainier, 2017; Kontarakis & Stainier, 2020). Here, we demonstrate that compensation by related genes buffers the effect of deleterious mutations in Chlamydomonas lipid biosynthesis genes, in a process that resembles transcriptional adaptation. Knockouts of *DGTT1* triggered upregulation of the functionally related and homologous *PDAT* gene regardless of the strain background, mutagenesis approach, or stress conditions (P or N deprivation), while knockout of *DGTT2* triggered upregulation of *PDAT*, *DGTT1* and *PGD1* (Fig. 1 and Fig. 2). In the *dgtt1*, *CC-dgtt1* and *dgtt2* mutant, the TAG-less phenotype found in respective knockdown lines (Liu et al., 2016) was not observed. Instead, these mutants produced more TAG than the control, likely due to transcriptional adaptation of homologous genes. This is unlikely the case for Arabidopsis mutants defective in TAG biosynthestic genes, where knockout of *DGAT1* and *PDAT1* resulted in a reduced TAG phenotype in oilseeds and vegetative tissues, respectively (Fan, Yan, & Xu, 2013; Zhang, Fan, Taylor, & Ohlrogge, 2009). Arabidopsis *DGAT1* and *PDAT1* are suggested to be functionally complementary genes essential for TAG biosynthesis and normal plant development (Fan et al., 2013; Zhang et al., 2009). Whether genetic compensation contributes to the observed phenotype in Arabidopsis mutants defective in TAG biosynthetic genes requires further study.

Recent advances in reverse genetic tools including CRISPR, TALEN, and ZFN genome editing methods have greatly accelerated progress in generating knockout mutants in traditional and new algal model systems, including Chlamydomonas (Greiner et al., 2017; Jiang et al., 2014; Park, Asbury, & Miller, 2020), *Nannochloropsis* (Poliner, Takeuchi, Du, Benning, & Farré, 2018; Wang et al., 2016), *Volvox carteri* (Ortega-Escalante, Jasper, & Miller, 2019), and *Ectocarpus* (Badis et al., 2021). Moreover, there is a genome-wide insertional mutant pool, the CLiP library (Xiaobo Li et al., 2019), for Chlamydomonas. Such tools and mutant libraries dramatically facilitate analysis of gene function. However, as we demonstrated here, genetic compensation by related genes might, in some cases, offset the effects of a null mutation, confounding genetic analysis. Likewise, previous efforts to screen and characterize TAG-less Chlamydomonas mutants from the CLiP library failed to identify any *DGTT* knockout mutants (XB Li, personal communication), possibly due to genetic compensation, while knockdown lines of *DGTTs* clearly showed a reduced-TAG phenotype (Liu et al., 2016). Therefore, when studying gene function using null mutants, preferably multiple mutant strains targeting the same gene should be obtained by a targeted genome editing method (e.g. CRISPR) or from the CLiP libarary (because some might activate genetic compensation while others might not), and these mutants should be carefully evaluated for their genotypic and phenotypic traits. If genetic compensation is observed in the mutants, alternative methods such as knockdown should be used to confirm the respective gene function. Ideally, deletion mutants should be analyzed as they will not trigger transcriptional adaptation through the nonsense-mediated mRNA decay pathway (El-Brolosy et al., 2019; Z. Ma et al., 2019), though to date a relatively small number of deletion mutants have been generated in Chlamydomonas due to current technical limitations.

In Chlamydomonas, biosynthesis of TAG has been implicated in the protection of the photosynthetic electron transport chain from over-reduction during stress conditions, and TAG accumulation is suggested to be essential for cells to survive under stress (Du et al., 2018; X. Li et al., 2012). Our initial experiments raised two interesting questions. First, does genetic compensation of TAG biosynthesis, as appears to occur in the *dgtt1* and *dgtt2* mutants, play an essential survival role under persistent stress conditions, and second, might genetic compensation occur for other mutated TAG biosynthetic genes? To address these questions, we first examined the *pdat* mutant. Unlike the *dgtt1* and *dgtt2* mutants, it has a similar TAG-less phenotype and TAG biosynthetic gene expression profiles when stressed as the knockdown lines (Fig. 3 and (Boyle et al., 2012; Yoon et al., 2012)), suggesting no genetic compensation in the *pdat* mutant. This outcome may be due to the poor function of *PDAT* in stress conditions, under which *DGTTs* are primarily responsible for TAG biosynthesis (Liu et al., 2016; Yoon et al., 2012). Consistent with this notion is a recent *PDAT* analysis in the green lineage showing that *PDAT* plays a less important role in algae than in plants (Falarz et al., 2020). Disruption of *AtPDAT1* in *A. thaliana* led to an over 50% decrease in oil accumulation in growing tissues and significant gametophytic and growth defects (Fan et al., 2013), whereas knockout of Chlamydomonas *PDAT* reduced TAG content by only 25% - 30%, and *pdat* cells had a similar growth rate as the wild type (Fig 3A) (Boyle et al., 2012).

To determine whether there might be compensation for the *PDAT* knockout when PDAT function is most essential, we evaluated the effect of the *PDAT* knockout in a Δ*vtc1* mutant, in which *PDAT* expression and function are likely critical for Δ*vtc1* cells to store excess energy as TAG under stress (Fig. 4). We expected that in the absence of compensation, the knockout of *PDAT* in the Δ*vtc1* background should result in a TAG-less phenotype. Intriguingly, the Δ*vtc1*/*pdat* mutant exhibited a 272.3% higher TAG content than the *VTC1* control, likely because of a large (12.8-fold) upregulation of *DGTT1* (Fig. 4). The Δ*vtc1*/*pdat* mutant remained viable under stress, as its cell density and dry weight increased in response to P deprivation (Fig. S4). So while no genetic compensation was observed in the *pdat* mutant, in the Δ*vtc1*/*pdat* double mutant, *DGTT1* was highly overexpressed, and more TAG was produced under P deprivation. Under stress conditions, these cells continue to capture light energy through photosynthesis, and if electron acceptors become overreduced due to the slow growth, cytotoxic reactive oxygen species are produced that lead to cell death (Y. Li, Sommerfeld, Chen, & Hu, 2008). TAG biosynthesis is likely essential for Δ*vtc1*/*pdat* cells to survive during stress because TAG is the most reduced electron sink in algae (Hu et al., 2008). In line with our findings, it was previously reported that a Chlamydomonas mutant defective for the MGDG-specific lipase PGD1 accumulated less TAG than the wild type while its cells lost viability under stress, suggesting TAG plays an essential role as an electron and energy sink under stress conditions, thereby preventing generation of reactive oxygen species (Du et al., 2018; X. Li et al., 2012). Together, these data suggest that under stress conditions, loss of TAG biosynthetic gene function can be buffered by upregulation of a related gene or genes through transcriptional adaptation in algae.

In summary, we have demonstrated that genetic compensation contributes to genetic robustness in microalgal TAG biosynthesis, maintaining TAG biosynthesis and redox homeostasis in knockout mutants under abiotic stress. This work demonstrates, to the best of our knowledge, the first reported example of genetic compensation in microalgae, implies the physiological relevance of genetic compensation in TAG biosynthesis under stress, and provides guidance for future genetic engineering and mutant characterization efforts. The exact mechanisms inducing genetic compensation in the Chlamydomonas mutants are not understood and warrant further investigation. Since transcriptional adaptation in other systems is triggered by mutant mRNA decay and involves Upf3a and COMPASS components (El-Brolosy et al., 2019; Z. Ma et al., 2019), testing the involvement of these components in algal genetic compensation is a logical next step.

## Materials and Methods

### Strains and growth conditions

*Chlamydomonas reinhardtii* LMJ.RY0402.223444, LMJ.RY0402.213587, LMJ.RY0402.223565 and their parent strain CC5325 (*cw15*; mt^−^) (Xiaobo Li et al., 2016), CC4502 (*pdat1-1*; mt^+^) and its parent strain CC4425 [D66] (*cw15*; mt^+^) (Boyle et al., 2012), CC3403 (*arg7*; *cw15*; mt^−^), Δ*vtc1* mutant strain CC5165 (Δ*vtc1*; mt^−^) and the *VTC1* rescue strain CC5166 (*ars76::VTC1*) (Aksoy et al., 2014) were obtained from the Chlamydomonas Resource Center (http://www.chlamycollection.org). CC3403 was used as the wild-type in CRISPR/Cas9 mutagenesis (Greiner et al., 2017). The Δ*vtc1*/*pdat* double mutant was derived from a cross between CC5165 and CC4502 (methods described below). All strains were maintained in Tris-acetate-phosphate (TAP) medium containing paramomycin as necessary. P deprivation was imposed by transferring *C. reinhardtii* cells to P-free TAP, in which potassium phosphate was replaced by 1.5 mM KCl. Ammonium chloride was omitted to generate N-free TAP. All batch cultures were grown under continuous light illumination at 60 μmol photons m^−2^ s^−1^, shaking at 150 rpm, room temperature.

### Gene expression measurement by RT-qPCR

Total RNA was extracted and purified using Trizol reagent (Invitrogen) and RNeasy mini kit (Qiagen). cDNA was synthesized using Protoscript II First Strand cDNA synthesis kit (NEB). The expression of target genes was determined by RT-qPCR in Applied Biosystems 7500 using the 2^−ΔΔCT^ method normalized by the expression of 18s rRNA. Details are described in the Supplemental Materials and Methods. The RT-qPCR analysis for each targeted gene was performed twice, with three biological replicates and three technical replicates for each sample. The Primer sequences used in RT-qPCR are listed in Table S2.

### Generation and screening of *Chlamydomonas reinhardtii* mutants

The CC*-dgtt1* mutant was generated by the CRISPR/Cas9 method developed by Greiner et al (Greiner et al., 2017). The parent strain of the *CC-dgtt1* mutant (CC3403), the SaCas9 expression vector (pHS_SaCas9) and the sgRNA cloning vector (pCrU6.4-SaCloning-aphVIII) were obtained from the Chlamydomonas Resource Center. In brief, a customized targeting vector that expressed sgRNA specifically targeting *DGTT1* was constructed in pCrU6.4-SaCloning-aphVIII. The SaCas9 expression vector and the customized targeting vector that co-transformed into CC3403 by electroporation (Park et al., 2020; Serobyan et al., 2020). The transformants were selected for paromomycin resistance and screened by PCR using oligonucleotides CrDGTT1F and CrDGTT1R (sequences in Table S2) for insertions at the *DGTT1* locus. The candidates yielding PCR products larger than the expected size for the wild type *DGTT1* region were sequenced. The detailed procedures are described in the Supplemental Materials and Methods.

The Δ*vtc1*/*pdat1* double mutant was generated by mating as described in Goodenough, 1976 (Goodenough, Hwang, & Martin, 1976) with some modifications (procedures described in the Supplemental Materials and Methods). The crossed candidates were selected for paramomycin resistance and subsequently screened by colony PCR for a co-occurrence with the deletion of the *VTC1* gene and the presence of the *pdat1-1* mutation (Boyle et al., 2012) in the genome. Primers CrVTC1-pF1 and CrVTC1-stopR1 (sequences in Table S2) were used for determination of the presence (yielding a 1148 bp PCR product) or the absence (yielding no PCR products) of the *VTC1* gene. Primers CrPDAT1-1-F2 and RIM5-2 (sequences in Table S2) (Boyle et al., 2012) were used to determine the presence (~ 1 kb PCR product) or the absence (no PCR product) of the *AphIII* insertion in the *PDAT* gene.

### TAG content measurement

Total lipid from *C. reinhardtii* cells was extracted using chloroform: methanol (2:1, v/v) as previously described (Yoon et al., 2012). The lipid extracts were dried under a gentle stream of nitrogen gas and re-dissolved in chloroform. Total lipid was resolved by thin-layer chromatography (TLC) on a silica gel 60 F_254_ plates (EMD Millipore) using hexane: t-butyl methyl ether: acetic acid (80:20:2, v/v/v) solvent mixture as a mobile phase to develop TAG. For visualization, the developed TLC plates were sprayed with 8% (w/v) H_3_PO_4_ solution containing 10% (w/v) copper (II) sulfate pentahydrate, and then charred at 180°C for 3 min. The relative intensity of signals corresponding to TAG can be measured using ImageJ software (NIH). The absolute quantification of TAG can be calculated by the sum of fatty acid fractions composed of TAG using gas chromatography-mass spectrometry (GC-MS) as previously described (Yantao Li et al., 2010; Ohlrogge & Browse, 1995). Details are described in the Supplemental Materials and Methods.

### Statistical analysis

Statistical analysis was performed by Student’s *t*-test with two-tailed distribution. *p* value ≤ 0.05 was the criteria to determine statistical significance.

## Acknowledgements

This work was supported in part by grants from the U.S. National Science Foundation (CBET-1511939) and DOE Office of Fossil Energy (FE-0031914) to YL, and by NSF 1332344 to SMM. We thank Drs. William Snell, Peeyush Ranjan, and Mayanka Awasthi at University of Maryland College park for their help with generating double knockout mutants, Dr. Arthur Grossman at Carnegie Institution for Science for providing the *Chlamydomonas vtc1* strains used in this study, Dr. Xiaobo Li at Westlake University for discussion of unpublished mutant screening data, and Drs. Fan Zhang and Russell Hill at the Institute of Marine and Environmental Technology for their help with polyP analysis. We thank Mr. Mohamed Mahmoud-Aly at Cairo University for his help with *Chlamydomonas* mutant characterization.

## References

Aksoy, M., Pootakham, W., & Grossman, A. R. (2014). Critical Function of a *Chlamydomonas reinhardtii* Putative Polyphosphate Polymerase Subunit during Nutrient Deprivation. The Plant Cell, 26(10), 4214–4229. doi: 10.1105/tpc.114.129270

Badis, Y., Scornet, D., Harada, M., Caillard, C., Godfroy, O., Raphalen, M., … Cock, J. M. (2021). Targeted CRISPR-Cas9-based gene knockouts in the model brown alga Ectocarpus. New Phytol. doi: 10.1111/nph.17525

Bagnato, C., Prados, M. B., Franchini, G. R., Scaglia, N., Miranda, S. E., & Beligni, M. V. (2017). Analysis of triglyceride synthesis unveils a green algal soluble diacylglycerol acyltransferase and provides clues to potential enzymatic components of the chloroplast pathway. BMC Genomics, 18(1), 223. doi: 10.1186/s12864-017-3602-0

Barabási, A. L., & Oltvai, Z. N. (2004). Network biology: understanding the cell’s functional organization. Nat Rev Genet, 5(2), 101–113. doi: 10.1038/nrg1272

Boyle, N. R., Page, M. D., Liu, B., Blaby, I. K., Casero, D., Kropat, J., … Merchant, S. S. (2012). Three acyltransferases and nitrogen-responsive regulator are implicated in nitrogen starvation-induced triacylglycerol accumulation in *Chlamydomonas*. Journal of Biological Chemistry, 287(19), 15811–15825. doi: 10.1074/jbc.M111.334052

Braun, N., Wyrzykowska, J., Muller, P., David, K., Couch, D., Perrot-Rechenmann, C., & Fleming, A. J. (2008). Conditional repression of AUXIN BINDING PROTEIN1 reveals that it coordinates cell division and cell expansion during postembryonic shoot development in Arabidopsis and tobacco. Plant Cell, 20(10), 2746–2762. doi: 10.1105/tpc.108.059048

Cases, S., Smith, S. J., Zheng, Y. W., Myers, H. M., Lear, S. R., Sande, E., … Farese, R. V., Jr. (1998). Identification of a gene encoding an acyl CoA:diacylglycerol acyltransferase, a key enzyme in triacylglycerol synthesis. Proc Natl Acad Sci U S A, 95(22), 13018–13023. doi: 10.1073/pnas.95.22.13018

Chen, X., Grandont, L., Li, H., Hauschild, R., Paque, S., Abuzeineh, A., … Friml, J. (2014). Inhibition of cell expansion by rapid ABP1-mediated auxin effect on microtubules. Nature, 516(7529), 90–93. doi: 10.1038/nature13889

Dahlqvist, A., Stahl, U., Lenman, M., Banas, A., Lee, M., Sandager, L., … Stymne, S. (2000). Phospholipid:diacylglycerol acyltransferase: an enzyme that catalyzes the acyl-CoA-independent formation of triacylglycerol in yeast and plants. Proc Natl Acad Sci U S A, 97(12), 6487–6492. doi: 10.1073/pnas.120067297

Du, Z. Y., Lucker, B. F., Zienkiewicz, K., Miller, T. E., Zienkiewicz, A., Sears, B. B., … Benning, C. (2018). Galactoglycerolipid Lipase PGD1 Is Involved in Thylakoid Membrane Remodeling in Response to Adverse Environmental Conditions in Chlamydomonas. Plant Cell, 30(2), 447–465. doi: 10.1105/tpc.17.00446

El-Brolosy, M. A., Kontarakis, Z., Rossi, A., Kuenne, C., Günther, S., Fukuda, N., … Stainier, D. Y. R. (2019). Genetic compensation triggered by mutant mRNA degradation. Nature, 568(7751), 193–197. doi: 10.1038/s41586-019-1064-z

El-Brolosy, M. A., & Stainier, D. Y. R. (2017). Genetic compensation: A phenomenon in search of mechanisms. PLoS genetics, 13(7), e1006780–e1006780. doi: 10.1371/journal.pgen.1006780

Falarz, L. J., Xu, Y., Caldo, K. M. P., Garroway, C. J., Singer, S. D., & Chen, G. (2020). Characterization of the diversification of phospholipid:diacylglycerol acyltransferases in the green lineage. Plant J, 103(6), 2025–2038. doi: 10.1111/tpj.14880

Fan, J., Yan, C., & Xu, C. (2013). Phospholipid:diacylglycerol acyltransferase-mediated triacylglycerol biosynthesis is crucial for protection against fatty acid-induced cell death in growing tissues of Arabidopsis. Plant J, 76(6), 930–942. doi: 10.1111/tpj.12343

Fedorov, Y., Anderson, E. M., Birmingham, A., Reynolds, A., Karpilow, J., Robinson, K., … Khvorova, A. (2006). Off-target effects by siRNA can induce toxic phenotype. RNA (New York, N.Y.), 12(7), 1188–1196. doi: 10.1261/rna.28106

Gao, Y., Zhang, Y., Zhang, D., Dai, X., Estelle, M., & Zhao, Y. (2015). Auxin binding protein 1 (ABP1) is not required for either auxin signaling or Arabidopsis development. Proc Natl Acad Sci U S A, 112(7), 2275–2280. doi: 10.1073/pnas.1500365112

Goodenough, U. W., Hwang, C., & Martin, H. (1976). Isolation and genetic analysis of mutant strains of Chlamydomonas reinhardi defective in gametic differentiation. Genetics, 82(2), 169–186.

Greiner, A., Kelterborn, S., Evers, H., Kreimer, G., Sizova, I., & Hegemann, P. (2017). Targeting of Photoreceptor Genes in Chlamydomonas reinhardtii via Zinc-Finger Nucleases and CRISPR/Cas9. Plant Cell, 29(10), 2498–2518. doi: 10.1105/tpc.17.00659

Hu, Q., Sommerfeld, M., Jarvis, E., Ghirardi, M., Posewitz, M., Seibert, M., & Darzins, A. (2008). Microalgal triacylglycerols as feedstocks for biofuel production: perspectives and advances. Plant J, 54, 621–639.

Iwai, M., Ikeda, K., Shimojima, M., & Ohta, H. (2014). Enhancement of extraplastidic oil synthesis in Chlamydomonas reinhardtii using a type-2 diacylglycerol acyltransferase with a phosphorus starvation-inducible promoter. Plant Biotechnology Journal, 12(6), 808–819. doi: 10.1111/pbi.12210

Jiang, W., Brueggeman, A. J., Horken, K. M., Plucinak, T. M., & Weeks, D. P. (2014). Successful transient expression of Cas9/sgRNA genes in Chlamydomonas reinhardtii. Eukaryotic Cell. doi: 10.1128/ec.00213-14

Kontarakis, Z., & Stainier, D. Y. R. (2020). Genetics in Light of Transcriptional Adaptation. Trends in Genetics, 36(12), 926–935. doi: 10.1016/j.tig.2020.08.008

Lardizabal, K. D., Mai, J. T., Wagner, N. W., Wyrick, A., Voelker, T., & Hawkins, D. J. (2001). DGAT2 is a new diacylglycerol acyltransferase gene family: purification, cloning, and expression in insect cells of two polypeptides from Mortierella ramanniana with diacylglycerol acyltransferase activity. J Biol Chem, 276(42), 38862–38869. doi: 10.1074/jbc.M106168200

Lenka, S. K., Carbonaro, N., Park, R., Miller, S. K., Thorpe, I., & Li, Y. T. (2016). Current advances in molecular, biochemical, and computational modeling analysis of microalgal triacylglycerol biosynthesis. Biotechnology Advances, 34, 1046–1063. doi: 10.1016/j.biotechadv.2016.06.004

Li, X., Moellering, E. R., Liu, B., Johnny, C., Fedewa, M., Sears, B. B., … Benning, C. (2012). A galactoglycerolipid lipase is required for triacylglycerol accumulation and survival following nitrogen deprivation in Chlamydomonas reinhardtii. Plant Cell., 24. doi: 10.1105/tpc.112.105106

Li, X., Patena, W., Fauser, F., Jinkerson, R. E., Saroussi, S., Meyer, M. T., … Jonikas, M. C. (2019). A genome-wide algal mutant library and functional screen identifies genes required for eukaryotic photosynthesis. Nature Genetics, 51(4), 627–635. doi: 10.1038/s41588-019-0370-6

Li, X., Zhang, R., Patena, W., Gang, S. S., Blum, S. R., Ivanova, N., … Jonikas, M. C. (2016). An Indexed, Mapped Mutant Library Enables Reverse Genetics Studies of Biological Processes in *Chlamydomonas reinhardtii*. The Plant Cell, 28(2), 367–387. doi: 10.1105/tpc.15.00465

Li, Y., Han, D., Hu, G., Dauvillee, D., Sommerfeld, M., Ball, S., & Hu, Q. (2010). *Chlamydomonas* starchless mutant defective in ADP-glucose pyrophosphorylase hyper-accumulates triacylglycerol. Metabolic Engineering, 12, 387–391. doi: DOI: 10.1016/j.ymben.2010.02.002

Li, Y., Sommerfeld, M., Chen, F., & Hu, Q. (2008). Consumption of oxygen by astaxanthin biosynthesis: A protective mechanism against oxidative stress in *Haematococcus pluvialis* (Chlorophyceae). J Plant Physiol. doi: 10.1016/j.jplph.2007.12.007

Liu, J., Han, D., Yoon, K., Hu, Q., & Li, Y. (2016). Characterization of type 2 diacylglycerol acyltransferases in Chlamydomonas reinhardtii reveals their distinct substrate specificities and functions in triacylglycerol biosynthesis. Plant Journal, 86, 3–19. doi: 10.1111/tpj.13143

Ma, Y., Creanga, A., Lum, L., & Beachy, P. A. (2006). Prevalence of off-target effects in Drosophila RNA interference screens. Nature, 443(7109), 359–363. doi: 10.1038/nature05179

Ma, Z., Zhu, P., Shi, H., Guo, L., Zhang, Q., Chen, Y., … Chen, J. (2019). PTC-bearing mRNA elicits a genetic compensation response via Upf3a and COMPASS components. Nature, 568(7751), 259–263. doi: 10.1038/s41586-019-1057-y

Miller, R., Wu, G., Deshpande, R. R., Vieler, A., Gartner, K., Li, X., … Benning, C. (2010). Changes in transcript abundance in Chlamydomonas reinhardtii following nitrogen deprivation predict diversion of metabolism. Plant Physiol, 154(4), 1737–1752. doi: 10.1104/pp.110.165159

Molnar, A., Bassett, A., Thuenemann, E., Schwach, F., Karkare, S., Ossowski, S., … Baulcombe, D. (2009). Highly specific gene silencing by artificial microRNAs in the unicellular alga *Chlamydomonas reinhardtii*. Plant Journal, 58(1), 165–174.

Oelkers, P., Cromley, D., Padamsee, M., Billheimer, J. T., & Sturley, S. L. (2002). The DGA1 gene determines a second triglyceride synthetic pathway in yeast. J Biol Chem, 277(11), 8877–8881. doi: 10.1074/jbc.M111646200

Ohlrogge, J., & Browse, J. (1995). Lipid biosynthesis. Plant Cell, 7(7), 957–970.

Ortega-Escalante, J. A., Jasper, R., & Miller, S. M. (2019). CRISPR/Cas9 mutagenesis in Volvox carteri. The Plant Journal, 97(4), 661–672. doi: https://doi.org/10.1111/tpj.14149

Park, R. V., Asbury, H., & Miller, S. M. (2020). Modification of a Chlamydomonas reinhardtii CRISPR/Cas9 transformation protocol for use with widely available electroporation equipment. MethodsX, 7, 100855. doi: https://doi.org/10.1016/j.mex.2020.100855

Plouviez, M., Fernández, E., Grossman, A. R., Sanz-Luque, E., Sells, M., Wheeler, D., & Guieysse, B. (2021). Responses of Chlamydomonas reinhardtii during the transition from P-deficient to P-sufficient growth (the P-overplus response): The roles of the vacuolar transport chaperones and polyphosphate synthesis. Journal of Phycology, 57(3), 988–1003. doi: https://doi.org/10.1111/jpy.13145

Poliner, E., Takeuchi, T., Du, Z. Y., Benning, C., & Farré, E. M. (2018). Nontransgenic Marker-Free Gene Disruption by an Episomal CRISPR System in the Oleaginous Microalga, Nannochloropsis oceanica CCMP1779. ACS Synth Biol, 7(4), 962–968. doi: 10.1021/acssynbio.7b00362

Rodriguez-Leal, D., Xu, C., Kwon, C.-T., Soyars, C., Demesa-Arevalo, E., Man, J., … Lippman, Z. B. (2019). Evolution of buffering in a genetic circuit controlling plant stem cell proliferation. Nature Genetics, 51(5), 786–792. doi: 10.1038/s41588-019-0389-8

Rohr, J., Sarkar, N., Balenger, S., Jeong, B. R., & Cerutti, H. (2004). Tandem inverted repeat system for selection of effective transgenic RNAi strains in Chlamydomonas. Plant Journal, 40(4), 611–621.

Rossi, A., Kontarakis, Z., Gerri, C., Nolte, H., Hölper, S., Krüger, M., & Stainier, D. Y. R. (2015). Genetic compensation induced by deleterious mutations but not gene knockdowns. Nature, 524(7564), 230–233. doi: 10.1038/nature14580

Sanz-Luque, E., Saroussi, S., Huang, W., Akkawi, N., & Grossman, A. R. (2020). Metabolic control of acclimation to nutrient deprivation dependent on polyphosphate synthesis. Sci Adv, 6(40). doi: 10.1126/sciadv.abb5351

Schmollinger, S., Chen, S., Strenkert, D., Hui, C., Ralle, M., & Merchant, S. S. (2021). Single-cell visualization and quantification of trace metals in Chlamydomonas lysosome-related organelles. Proc Natl Acad Sci U S A, 118(16). doi: 10.1073/pnas.2026811118

Schroda, M. (2006). RNA silencing in Chlamydomonas: mechanisms and tools. Current Genetics, 49(2), 69–84. doi: 10.1007/s00294-005-0042-1

Serobyan, V., Kontarakis, Z., El-Brolosy, M. A., Welker, J. M., Tolstenkov, O., Saadeldein, A. M., … Stainier, D. Y. R. (2020). Transcriptional adaptation in Caenorhabditis elegans. eLife, 9, e50014. doi: 10.7554/eLife.50014

Tautz, D. (1992). Redundancies, development and the flow of information. Bioessays, 14(4), 263–266. doi: 10.1002/bies.950140410

Teng, X., Dayhoff-Brannigan, M., Cheng, W.-C., Gilbert, Catherine E., Sing, Cierra N., Diny, Nicola L., … Hardwick, J. M. (2013). Genome-wide Consequences of Deleting Any Single Gene. Molecular Cell, 52(4), 485–494. doi: https://doi.org/10.1016/j.molcel.2013.09.026

Wang, Q., Lu, Y., Xin, Y., Wei, L., Huang, S., & Xu, J. (2016). Genome editing of model oleaginous microalgae Nannochloropsis spp. by CRISPR/Cas9. Plant J, 88. doi: 10.1111/tpj.13307

Yoon, K., Han, D., Li, Y., Sommerfeld, M., & Hu, Q. (2012). Phospholipid:diacylglycerol acyltransferase is a multifunctional enzyme involved in membrane lipid turnover and degradation while synthesizing triacylglycerol in the unicellular green microalga *Chlamydomonas reinhardtii*. Plant Cell, 24, 3708–3724. doi: 10.1105/tpc.112.100701

Zhang, M., Fan, J., Taylor, D. C., & Ohlrogge, J. B. (2009). DGAT1 and PDAT1 acyltransferases have overlapping functions in Arabidopsis triacylglycerol biosynthesis and are essential for normal pollen and seed development. The Plant Cell, 21(12), 3885–3901. doi: 10.1105/tpc.109.071795

