## Supplemental material for "Genetic compensation of triacylglycerol biosynthesis in the green microalga *Chlamydomonas reinhardtii*"

**Supplemented Materials and Methods**

**Setting up of P stress conditions**

Seed cultures were grown in TAP medium until late exponential or early stationary phases and then inoculated into fresh medium with 1 x 10^6^ cells/ml initial cell density. Cell density was measured using a hemocytometer (Hausser Scientific). To impose P deprivation, the inoculum was washed with P-free TAP (TAP-P) medium twice before inoculation. All batch cultures were grown under continuous light illumination at 60 μmol photons m^-2^ s^-1^, shaking at 150 rpm, and room temperature.

**RNA isolation and quantitative real-time PCR**

Total RNA was isolated from *C. reinhardtii* cells using Trizol reagent (Invitrogen) and quantified using NanoDrop 2000c spectrophotometer (Thermo Scientific). One μg RNA was converted to cDNA using Protoscript II First Strand cDNA synthesis kit (NEB). The expression of target genes was determined by quantitative real-time PCR (qRT-PCR) using Applied Biosystems 7500. Each qRT-PCR reaction contains 25 ng cDNA, 400 nM of each primer pair and 10 μl of SYBR Green PCR Master Mix (Invitrogen) to a final volume of 20 μl. PCR cycling conditions consisted of an initial polymerase activation step at 95^°^C for 30 s followed by 40 cycles at 95^°^C for 5 s and 60^°^C for 30 s. The expression of 18s rRNA was used as the reference. The relative expression level was measured using the 2^-ΔΔCT^ method. The experiment was performed twice, with three biological replicates and three technical replicates for each sample. The Primer sequences used in qRT-PCR are listed in Table S2.

**Generation of *Chlamydomonas reinhardtii* mutants by CRIPR/Cas9 mutagenesis**

The CC*-dgtt1* mutant was generated by the CRISPR/Cas9 method developed by Greiner et al (17). CC3403, the parent strain of the *CC-dgtt1* mutant, the SaCas9 expression vector (pHS_SaCas9, catalog # pPH187) and the sgRNA cloning vector (pCrU6.4-SaCloning-aphVIII, catalog # pPH339) that expressed customized sgRNA targeting *DGTT1* were obtained from the Chlamydomonas Resource Center ([http://www.chlamycollection.org](http://www.chlamycollection.org/)). The customized targeting vector was made by annealing oligonucleotides (5’- acttGGTCTTAATCAGGCGGGCGCCG-3’ and 5’- aaacCGGCGCCCGCCTGATTAAGACC-3’) and ligating into *Esp3I*-digested pCrU6.4-SaCloning-aphVIII. This targeting vector and pHS_SaCas9 were co-transformed by electroporation into CC3403 (9, 41). Paromomycin-resistant transformants were selected and screened by PCR for insertions at the *DGTT1* locus as follows. Paromomycin-resistant colonies were transferred to wells of a 96-well plate each containing 180 μl of TAP containing 100 μg/mL _L_-arginine and 10 µg/mL paromomycin. The plates were grown under room light (2 - 4 µmol photons m^-2^ s^-1^ for 10 - 12 hours per day) at room temperature for 7 - 12 days. 40 μL from each well was transferred to a PCR tube and centrifuged for 10 min at 2000 *g* at room temperature. The pellet was resuspended in 20 μl of Dilution Buffer from the Phire Plant Direct PCR Master Mix Kit (ThermoFisher Scientific) and incubated for 5 min at room temperature. Samples were centrifuged for 10 min at 2000 *g* at room temperature, then 15 μL of supernatant was removed to another PCR tube with 60 μL distilled water to dilute the sample prior to PCR. The PCR reaction consisted of 2.5 μL distilled water, 5 μl 2x Phire Plant Direct PCR Master Mix, 1 μL forward primer (10 μM), 1 μL reverse primer (10 μM), and 0.5 μL DNA extract solution. Reactions included oligonucleotides CrDGTT1F (5’-CTCTGCTCATCGGCACATTG-3’) and CrDGTT1R (5’-ATATGCCACTTGCGGAAGGT-3’) in a Bio-Rad T100 Thermal Cycler (Hercules, CA, USA) programmed to 98°C for 5 min followed by 35 cycles of 98°C for 5 s, 65°C for 5 s, 72°C for 60 s, and a final extension step at 72°C for 30 s. Reaction products larger than that expected for the wild type *DGTT1* were sequenced to identify the *dgtt1* mutant.

**Generation of *Chlamydomonas reinhardtii* mutants by crossing**

Cultures of *Chlamydomonas* were grown in M medium under light dark cycle (13 h light: 11 h dark) until reaching the exponential phase and then switched to N-free M medium under continuous light at 170 rpm shaking for overnight to induce gamate formation. Gamate cells were harvested by centrifugation at 3000 rpm and room temperature for 5 min and washed with N-free M medium twice before resuspended in fresh N-free medium. Equal number of gamate cells of the opposite mating types were mixed together with addition of 150 mM db-cAMP and incubated under light for up to 2 h. 200 µL portion of gamates mixture were spreaded on a TAP medium plate containing 2% agar at every 30 min of the 2 h incubation. The plates were exposed to light for overnight and then place in the dark for 5 – 7 days to allow zygote formation. At the end of the incubation period, poorly adhered vegetative cells were gently scraped off from the plate surface by a razor blade and killed by the vapor of chloroform (30 – 60 sec exposure). The plates were exposed to light for overnight to allow zogotes to germinate before incubated under light dark cycles (13 h light: 11 h dark) to allow colonies to grow. Once zygotic colonies became visible, they were picked and resuspended in 200 µL TAP medium, and subsequently plated on a TAP medium containing 1.5% agar and 15 μg/mL paramomycin. The plates were incubated under continuous light to allow the paramomycin-resistent mutant candidates to grow.

**Fatty acid analysis by GC-MS**

TAG fractions in the lipid extracts on a developed TLC plate were visualized by I_2_ vapor and isolated by scrapping off the silica gel from the TLC plate. Fatty acids of the isolated TAG fractions were converted to fatty acid methyl esters (FAMEs) with 1% H_2_SO_4_ in methanol at 85^o^C for 1.5 h, dissolved in hexane, and then profiled using TSQ 8000 Triple GC-MS System (Thermo Scientific). FAME standards (Sigma) and heptadecanoic acid (C17:0) (Sigma) were used as the external and internal standards, respectively, for fatty acid analysis. The quantification for TAG was calculated by the sum of fatty acid amount based on GC-MS analysis.

Table S1. Pair-wise sequence similarity between TAG biosynthetic genes calculated by LALIGN program

| Gene 1 | Gene 2 | Identity matrix (genome) | Identity matrix (transcript) |
| --- | --- | --- | --- |
| *PDAT* | *DGTT1* | 50.9% | 50.5% |
| *PDAT* | *DGTT2* | 49.8% | 50.7% |
| *PDAT* | *DGTT3* | 49.2% | 52.8% |
| *DGTT1* | *DGTT2* | 50.5% | 51.4% |
| *DGTT1* | *DGTT3* | 49.0% | 51.6% |
| *DGTT2* | *DGTT3* | 50.6% | 59.1% |

Table S2. Primer sequences used in this study

| Primer name | Sequence (5’ to 3’) |
| --- | --- |
| CrDGTT1F | CTCTGCTCATCGGCACATTG |
| CrDGTT1R | ATATGCCACTTGCGGAAGGT |
| CrDGTT2F | CACCGACAAATGTGCGAATT |
| CrDGTT2R | CACATGCATCCAGCCACAGT |
| CrDGTT2F2 | TGCTGGACTCGAAGAAGA |
| CrDGTT2R2 | GATGTCAAAGCCGGGAAA |
| CrDGTT3F | ACCTCGCACTTGACCCTGAA |
| CrDGTT3R | TCATGAAGCCTACATAAATCGACATC |
| CrDGTT3F2 | CCCGAATGTTCGCATCTAC |
| CrDGTT3R2 | CCACCAGCAGCATGAAATA |
| CrPDATF | CTCCCAGTCTCAGCAGC |
| CrPDATR | GTAGGCATTGAGGCGAGG |
| CrPDATF2 | CGGGAGCCATGAGAGTG |
| CrPDATR2 | GATCACATCCCGCAGCAC |
| CrPGD1F | AGCCAGCTATTGTCGCACTT |
| CrPGD1R | CAAGAAATCCGCTGACATCC |
| Cr18SrRNAF | ACGAGACCTCAGCCTGCTAAAT |
| Cr18SrRNAR | TTATCGCCTCATACTTCCATTGG |
| CrVTC1-pF1 | TGCCATACAGATTTATCCGATG |
| CrVTC1-stopR1 | GAAAAGGTAGTCAATGACGGCG |
| CrPDAT1-1-F2 | GGTCACACGGCTGGAC |
| RIM5-2 | GCTGGCACGAGTACGGGTTG |

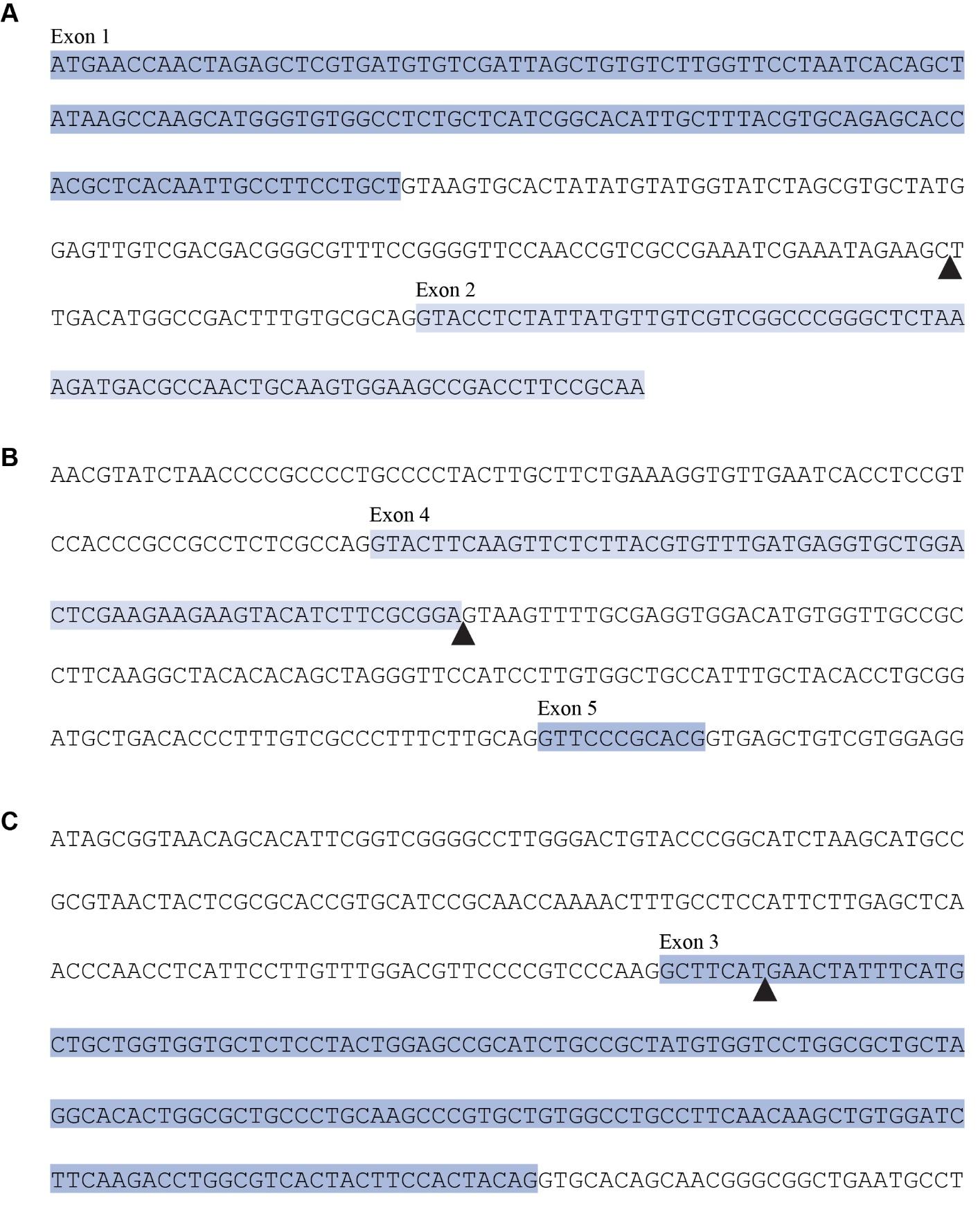

Fig S1. Locations of the CIB1 cassette (black triangle) insertion in the (A) *DGTT1*, (B) *DGTT2*, and (C) *DGTT3* genes in the *dgtt1* (LMJ.RY0402.223444), the *dgtt2* (LMJ.RY0402.213587), and the *dgtt3* (LMJ.RY0402.223565) strains.

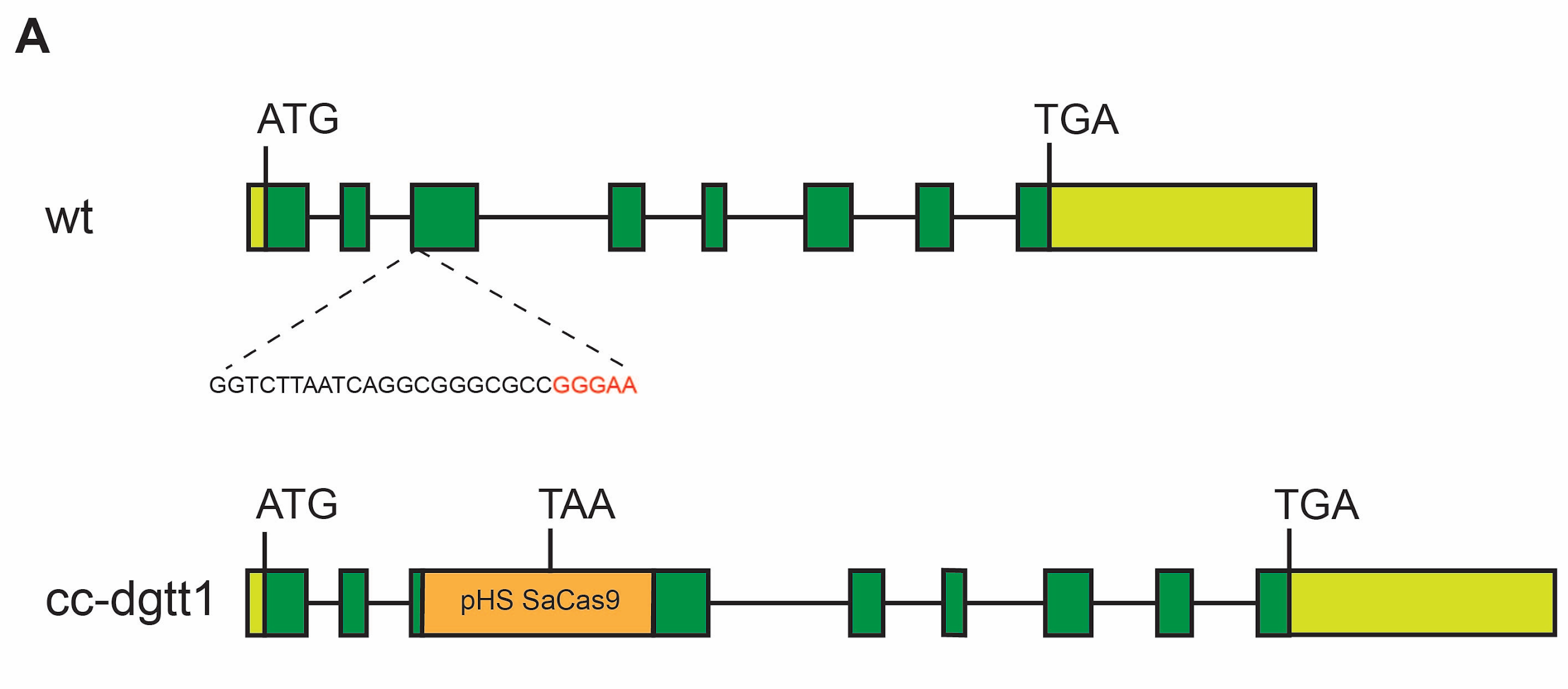

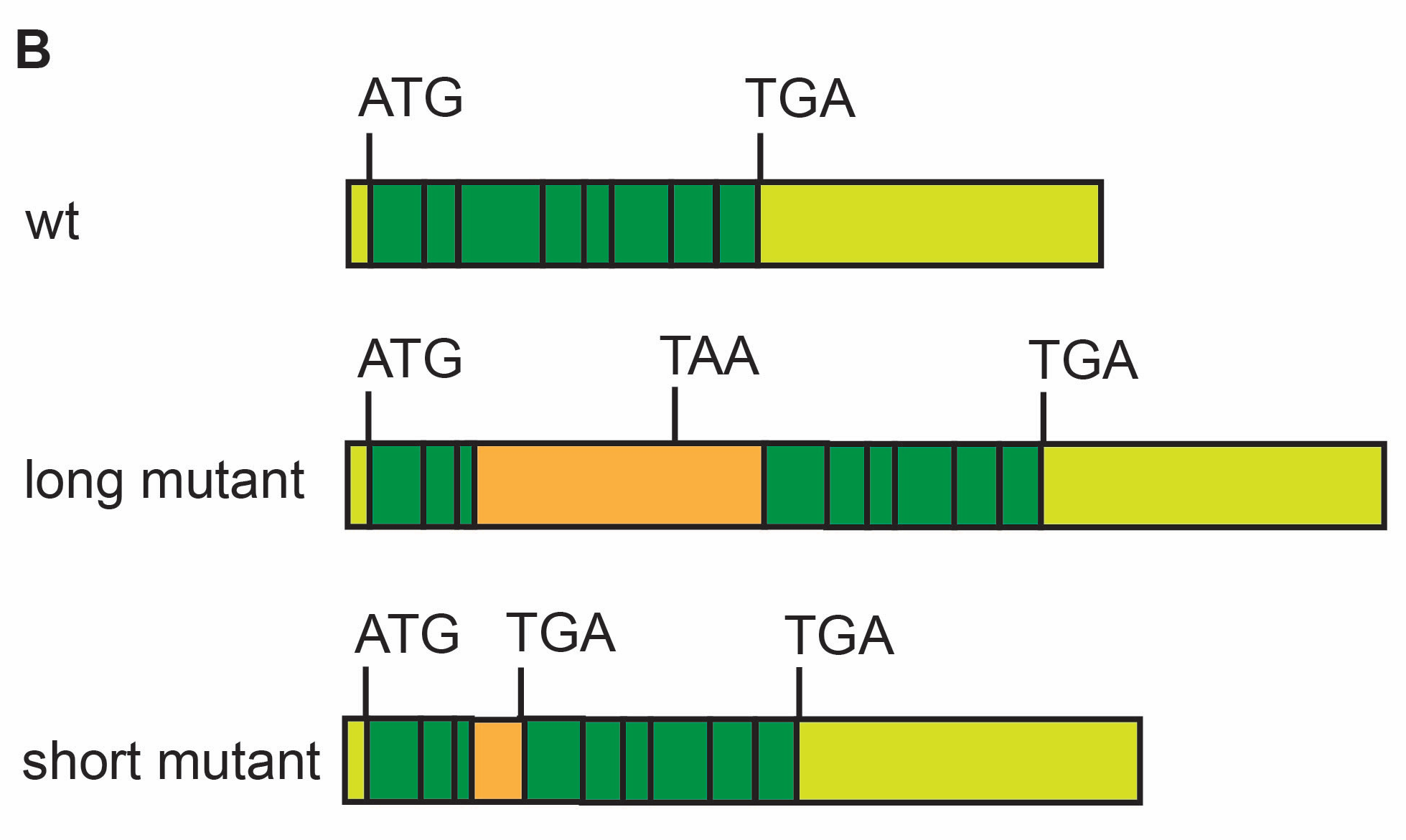

Fig. S2. Location of the inserted pHS SaCas9 plasmid fragment in *DGTT1* CRISPR/Cas9 mutant *CC-dgtt1*, and structure of *DGTT1* transcripts generated by the mutant. (A) Schematics showing wild type *DGTT1* gene model, with position of region targeted for SaCas9 mutagenesis indicated near beginning of exon 3(top) and gene model of mutant with 1229-bp fragment of SaCas9 plasmid inserted at targeted region. (B) Schematics showing structures of wild type *DGTT1* transcript (top), and the long aberrant (middle) and short aberrant (bottom) transcripts that were the only *DGTT1* transcripts detected in the mutant by RT-PCR. Both aberrant transcripts contain premature stop codons and code for non-functional DGTT1 proteins with only ~20% of the conserved diacylglycerol acyltransferase domain. Light green boxes, 5’ and 3’ UTRs; dark green boxes, exons; orange boxes, fragments of pHS Cas9 sequence.

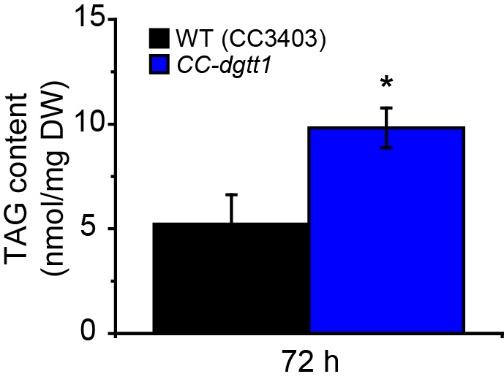

Fig S3. TAG content of the *CC-dgtt1* mutant (blue) and the parent strain CC3403 (black) grown was measured at 72 h under N deprivation. Data represents mean ± standard deviation (SD) from three biological repeats. An asterisk indicates significance by Students *t*-test (*p* value ≤ 0.05).

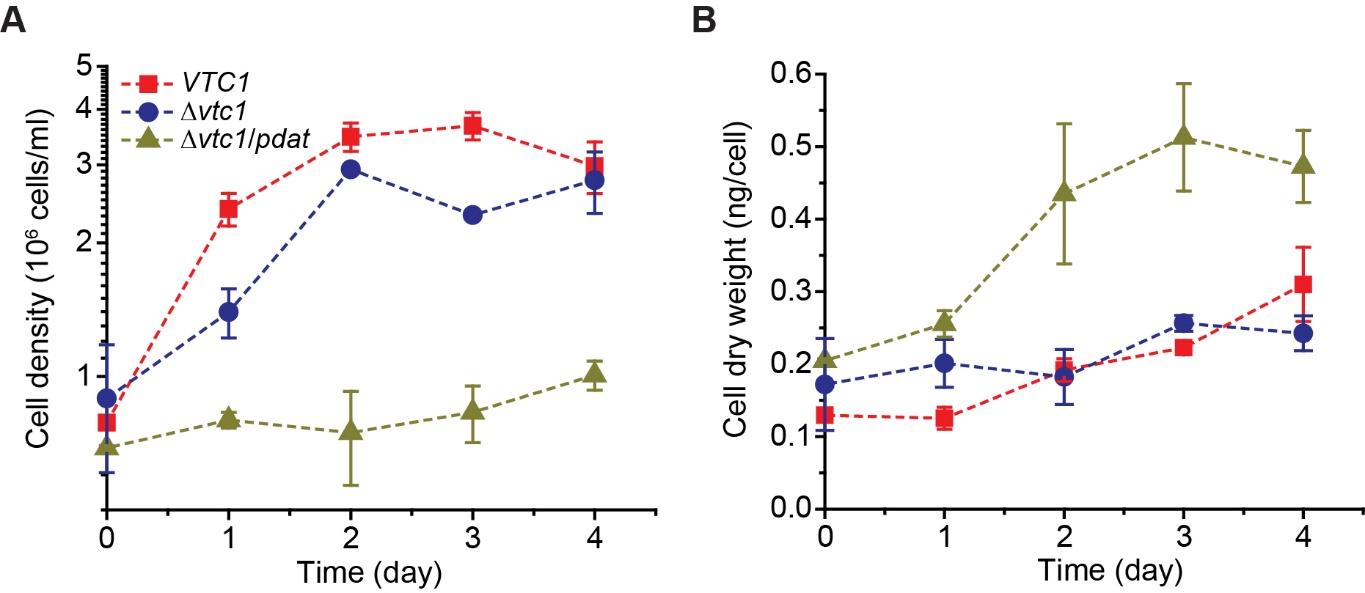

Fig S4. Growth curves in cell density (A) and cell dry weight (B) of batch cultures of *C. reinhardtii* *VTC1*, Δ*vtc1* and Δ*vtc1*/*pdat1* strains under P deprivation. Values are mean ± standard deviation (SD) from three biological repeats.
